# Interaction between spike protein of SARS-CoV-2 and human virus receptor ACE2 using two-color fluorescence cross-correlation spectroscopy

**DOI:** 10.1101/2021.08.18.456769

**Authors:** Ai Fujimoto, Lyu Yidan, Masataka Kinjo, Akira Kitamura

## Abstract

Infection with severe acute respiratory syndrome coronavirus-2 (SARS-CoV-2), the cause of coronavirus disease 2019 (COVID-19), is initiated by the interaction between a receptor protein, angiotensin-converting enzyme type 2 (ACE2) on the cell surface, and the viral spike (S) protein. This interaction is similar to the mechanism in SARS-CoV, a close relative of SARS-CoV-2, which was identified in 2003. Drugs and antibodies that inhibit the interaction between ACE2 and S proteins could be key therapeutic methods for preventing viral infection and replication in COVID-19. Here, we demonstrate the interaction between human ACE2 and a fragment of the S protein (S1 subunit) derived from SARS-CoV-2 and SARS-CoV using two-color fluorescence cross-correlation spectroscopy (FCCS), which can detect the interaction of fluorescently labeled proteins. The S1 subunit of SARS-CoV-2 interacted in solution with soluble ACE2, which lacks a transmembrane region, more strongly than that of SARS-CoV. Furthermore, one-to-one stoichiometry of the two proteins during the interaction was indicated. Thus, we propose that this FCCS-based interaction detection system can be used to analyze the interaction strengths of various mutants of the S1 subunit that have evolved during the worldwide pandemic, and also offers the opportunity to screen and evaluate the performance of drugs and antibodies that inhibit the interaction.

## 1. Introduction

Coronavirus disease 2019 (COVID-19) is caused by severe acute respiratory syndrome coronavirus-2 (SARS-CoV-2), a positive-strand RNA virus. The worldwide pandemic caused by this virus, which began in early 2020, has severely restricted economic and social activities around the world. The genome of SARS-CoV-2 shares approximately 80% identity with that of SARS-CoV, the virus causing SARS, which was identified in 2003 [1,2]. In both cases, spike glycoproteins (S proteins) on the surface of the virus recognize human cell receptors and mediate membrane fusion between the virus and human cells [3–5]. The S protein is cleaved into the S1 and S2 subunits during viral infection. The S1 subunit contains the receptor-binding domain (RBD) [6,7].

Human angiotensin-converting enzyme type 2 (hACE2) is the receptor of SARS-CoV on the human cell surface; as such, hACE2 was promptly identified as the receptor of SARS-CoV-2 [1,6,7]. ACE2 is a type I membrane protein containing a single transmembrane helix and a ~40-residue cytoplasmic segment that is expressed in the lungs, heart, kidneys, and intestine [8,9]. The S protein of SARS-CoV-2 interacts with hACE2 with a dissociation constant (*K*_d_) of several tens of nanomolars, based on surface plasmon resonance (SPR) analysis; this is ~10 to 20-fold higher than that of SARS-CoV [5,10,11]. Neutralizing antibodies, which bind to the S protein and prevent its interaction with ACE2, are useful for preventing viral infection [12–14]. Vaccines, which ideally produce neutralizing antibodies against the S protein, are an extremely useful way to produce immunity for the prevention of SARS-CoV-2 [15–17]. Current treatment strategies for COVID-19 are mainly focused on chemical medicines, including various neutralizing antibodies [13,18,19]. Thus, it is important to characterize inhibitory chemicals, such as antibodies and drugs, to prevent SARS-CoV-2 infection [20].

Fluorescence correlation spectroscopy (FCS) and its advanced method using two-color fluorescence, fluorescence cross-correlation spectroscopy (FCCS), have been used for biomolecular interactions with single-molecule sensitivity [21–25]. The most important feature of FCS/FCCS is a solution measurement system that does not use a solid-liquid interface, as in SPR. Based on these characteristics, screening for interaction inhibitors has been conducted. This approach also benefits from the requirement of only a small amount of liquid and a short measurement time [23]. These features suggest that FCCS has potential as a screening method for interaction inhibition; however, the interaction between S protein and ACE2 has not been demonstrated using FCCS. In this study, we analyzed the interaction between the fluorescent protein-tagged recombinant ACE2 and the S1 subunit expressed in mouse culture cells using FCCS.

## Methods

### Preparation of plasmid DNA

Human ACE2 (hACE2) cDNA was obtained from Addgene (#1786; Watertown, MA, USA). To construct an expression plasmid, cytoplasmic region- and transmembrane-lacking hACE2 tagged with a monomeric variant of eGFP carrying the A206K mutation (hACE2-eGFP), the fragment encoding hACE2, was inserted into pmeGFP-N1 with NheI and EcoRI. To reconstruct the linker region between hACE2 and eGFP, a double-stranded synthetic oligonucleotide (5′-gaattcttttgtgggatggagtaccgactggagtccatatgcagacggaccggt-3′) was inserted into the plasmid with EcoRI and AgeI. A polyhistidine tag was inserted into the C-terminus of eGFP. The whole sequence of the coding region of polyhistidine-tagged hACE2-eGFP is shown in Supplemental Figure. A synthetic oligonucleotide for the ER-targeting signal peptide was inserted into pmCherry-N1 (pER-mCherry) with NheI and AgeI. The DNA fragments of the spike protein 1 subunit (306–575 region) of SARS-CoV (#NC_004713) and SARS-CoV-2 (#NC_045512), including a polyhistidine tag at the C-terminus (S1 and S1-2, respectively) were synthesized by Eurofins Genomics GmbH (München, Germany). The fragments were inserted into pER-mCherry-N1 with XhoI and BamHI (pER-mCherry-S1 and pER-mCherry-S1-2, respectively). The whole sequences of the coding regions of pER-mCherry-S1 and pER-mCherry-S1-2 are presented in Supplemental Figure. The expression plasmid for soluble secreted yellow fluorescent protein (ssYFP)-tagged BiP (ssYFP-BiP), an ER-resident ortholog of heat shock protein 70 kDa, as an ER marker was kindly provided by Dr. Ikuo Wada (Fukushima Medical University, Fukushima, Japan).

### Cell culture and transfection

Mouse neuroblastoma Neuro2a cells were maintained as previously reported [26]. The cells (2.0 × 10^5^) were grown in 15 cm dishes (#TR4003, Nippon Gene, Tokyo, Japan) for 16 h before transfection. Plasmids encoding hACE2-eGFP, ER-mCherry-S1, or ER-mCherry-S1-2 (16 μg) and sonicated salmon sperm DNA (14 μg) were transfected into the cells with 240 μL of polyethyleneimine (#43896, Alfa Aesar, Ward Hill, MA, USA). After incubation of the cells for 24 h, subsequent experiments were performed. To confirm the intracellular localization of transiently expressed hACE2-eGFP, ER-mCherry-S1, and ER-mCherry-S1-2, these plasmids (1.0 μg) and plasmids encoding fluorescently visualized ER-marker proteins (ssYFP-BiP or ER-mCherry) (0.2 μg) were transfected into Neuro2a cells using Lipofectamine 2000 (Thermo Fisher) in a 35 mm glass base dish (IWAKI, Shizuoka, Japan). After incubation of the cells for 24 h, the cells were observed using confocal microscopy.

### Confocal microscopy

Neuro2a cells expressing hACE2-eGFP + ER-mCherry, ssYFP-BiP + ER-mCherry-S1, and ssYFP-BiP + ER-mCherry-S1-2 were grown in a 35 mm glassbase dish (IWAKI). Images of cells were acquired using a confocal laser scanning microscope (LSM510 META, Carl Zeiss) with a C-Apochromat 40×/1.2 NA Korr UV-VIS-IR water immersion objective (Carl Zeiss). The eGFP and ssYFP were excited at 488 nm; mCherry was excited at 594 nm, excited light and reflected fluorescence were separated using a beam splitter (HFT488/594). Fluorescence was selected using 500–530 nm bandpass (BP500– 530) and 615 nm long pass (LP615) filters. Pinholes were set at 71 μm for eGFP or ssYFP, and 83 μm for mCherry. For observation of lysosome localization, mCherry-tagged protein-expressing Neuro2a cells were incubated in a medium containing 0.5 μM Lysotrascker green DND-26 (Thermo Fisher) for 30 min before image acquisition.

### Protein purification

Neuro2a cells transiently expressing polyhistidine-tagged hACE2-eGFP, ER-mCherry-S1, ER-mCherry-S1-2, eGFP, and mCherry were lysed in 50 mM HEPES-KOH (pH 7.5), 150 mM NaCl, 1% Noidet P-40 (NP-40), and protease inhibitor cocktail (Sigma-Aldrich). The supernatants recovered through centrifugation (20,400 × *g*, 10 min, 4°C) were mixed with Ni-NTA agarose beads (Fujifilm Wako Pure Chemical Corp., Osaka, Japan) equilibrated in a buffer containing 50 mM HEPES-KOH (pH 7.5), 150 mM NaCl, 0.2% NP-40, and 10 mM imidazole. The beads were washed three times in wash buffer containing 50 mM HEPES-KOH (pH 7.5), 150 mM NaCl, 0.2% NP-40, and 20 mM imidazole. The proteins were eluted in a buffer containing 50 mM HEPES-KOH (pH 7.5), 150 mM NaCl, 0.2% NP-40, and 250 mM imidazole, followed by buffer exchange (50 mM HEPES-KOH, pH 7.5 and 150 mM NaCl) and concentration processes performed using a dialysis Amicon Ultra 10 K (Merck Millipore, Darmstadt, Germany).

### SDS-PAGE and western blotting

A portion of purified sample was mixed with SDS-PAGE sample buffer containing 50 mM dithiothreitol and boiled. Samples were applied to a 12.5% polyacrylamide gel and subjected to electrophoresis in SDS-containing buffer. Protein-transferred PVDF membranes (GE Healthcare Life Science) were blocked with 5% skim milk in PBS-T. The antibodies used were anti-GFP HRP-DirecT (#598-7, MBL, Nagano, Japan), anti-polyhistidine-tag (#PM032, MBL), and anti-mCherry (#Z2496N, TaKaRa, Shiga, Japan).

### Fluorescence cross-correlation spectroscopy (FCCS)

FCCS measurements were performed using a ConfoCor 3 system combined with an LSM 510 META (Carl Zeiss, Jena, Germany) through a C-Apochromat 40×/1.2 NA Korr UV-VIS-IR water-immersion objective (Carl Zeiss). The confocal pinhole diameter was adjusted to 73 μm. The eGFP and mCherry were excited at 488 and 594 nm, respectively, and emission signals were detected using a 505–540 nm band-pass filter for eGFP and a 615–680 nm band-pass filter for mCherry after the short and long wavelengths were divided through a 600 nm dichroic mirror. Measurements were performed in a cover glass chamber (#155411, Thermo Fisher Scientific, Waltham, MA, USA). The obtained fluorescence autocorrelation functions for eGFP and mCherry, *G*(τ)s, from which the lag time (τ) was analyzed using a one-component three-dimensional diffusion model including the one-component triplet state, is given by Eq. 1:

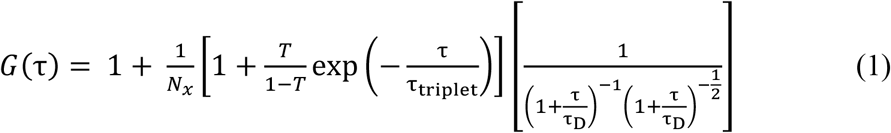

where τ_D_ is the diffusion time of molecules, *N_x_* is the average number of fluorescent molecules (eGFP-tagged and mCherry-tagged proteins are shown as *N*_g_ and *N*_r_, respectively) in the detection volume defined by the beam waist *w_xy_* and the axial radius *w*_z_, *s* is a structure parameter representing the ratio of *w_xy_* and *w*_z_, *T* is the blinking fraction, and τ_triplet_ is the blinking time. The *G*(τ) values of the samples were measured for 100 s. Following pinhole adjustment, the structure parameter was determined and fixed before measurements using a 10 nM ATTO488 and 10 nM Alexa Fluor 594 solution as a standard. Counts per molecule (CPM) were calculated as the mean fluorescence intensity of GFP divided by *N*_g_. The cross-correlation functions (CCFs) of the two fluorescent proteins were measured after mixing 10 nM eGFP-tagged hACE2 and 300 nM mCherry-tagged S1 subunit for 1 min; then, the CCFs were fitted using Eq. 1 with *T* = 0.

The relative cross-correlation amplitude was calculated using Eq. 2, as previously reported [26]:

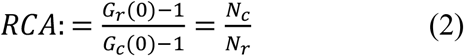

where *G*_r_(0) is the autocorrelation function of the red channel with mCherry at *t* = 0, *G*_c_(0) is the cross-correlation function between green with eGFP and red channels at *t* = 0, and *N*_c_ is the average number of interacting molecules.

### Statistical analysis

One-way analysis of variance (ANOVA) with post-hoc Tukeys significant difference test was performed using Origin Pro 2021 (OriginLab Corp., Northampton, MA).

## Results

Murine neuroblastoma Neuro2a (N2a) cells are known to show high protein expression owing to their high plasmid DNA transfection efficiency. Since N2a cells have high protein expression efficiency, N2a cells were selected as a cell line for SARS-CoV-associated protein expression. To establish a strategy for protein purification from N2a cells, subcellular localization of transmembrane-lacking hACE2, S1, and S1-2 subunits tagged with fluorescent proteins in N2a cells was confirmed using confocal fluorescence microscopy. The hACE2-eGFP (ACE2), ER-mCherry-S1 (S1), and ER-mCherry-S1-2 (S1-2) were observed inside the subcellular structures, including the ER (Figure 1A). Some ER-mCherry foci were colocalized with lysotracker green (LysoG Figure 1B). Since monomeric red fluorescent protein (mRFP) and its variants, such as mCherry, are often observed in lysosomes, mainly because of their low p*K*_a_ [27,28], these results suggest that ACE2, S1, and S1-2 are efficiently expressed and localized in various secretory pathways, including the ER and lysosome.

**Figure 1.**
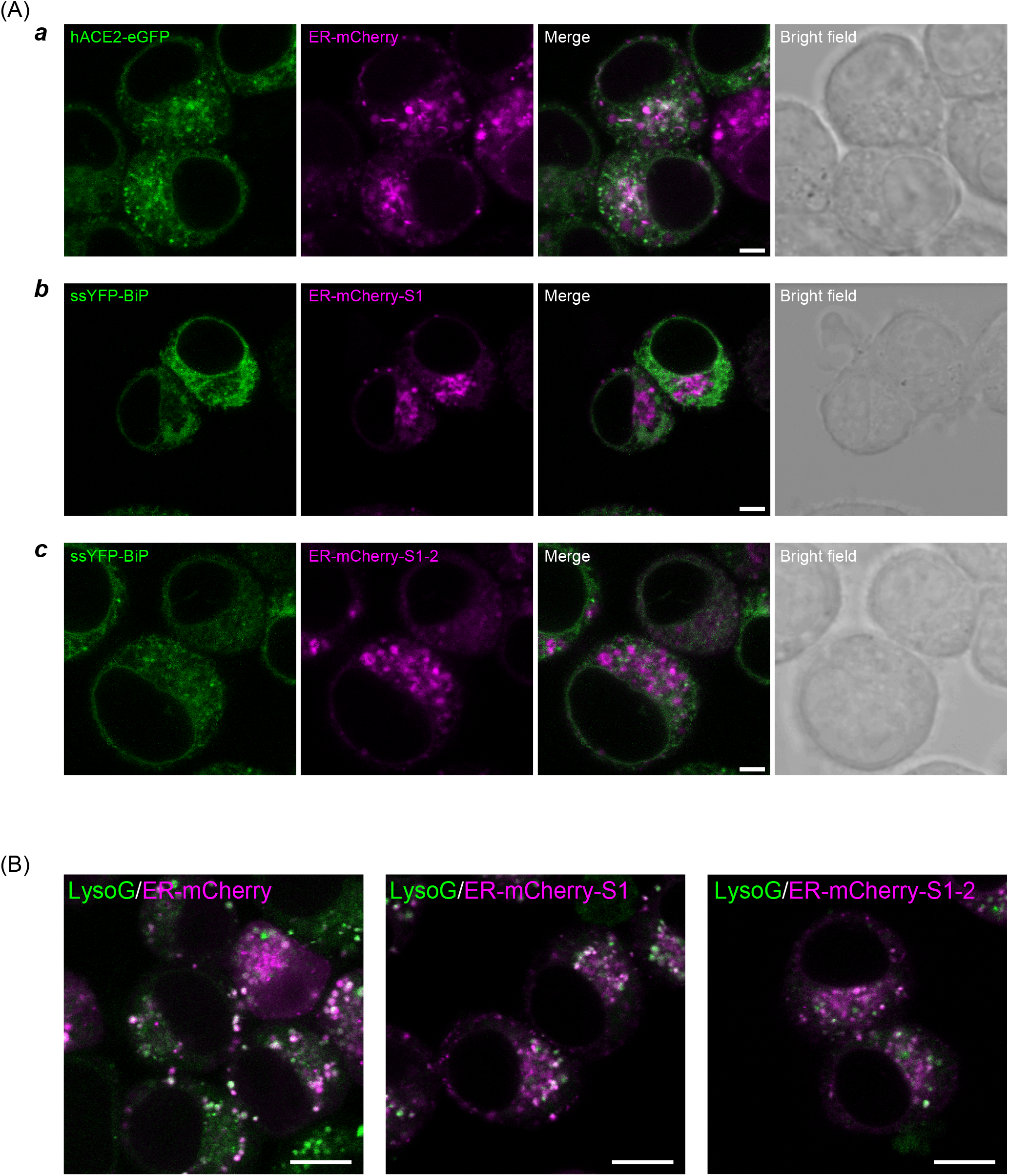
Confocal fluorescent images of murine neuroblastoma Neuro2a cells expressing eGFP-tagged hACE2 and mCherry-tagged S1 proteins. (A) Fluorescence images of Neuro2a cells expressing hACE2-eGFP, ER-mCherry-S1, or ER-mCherry-S1-2. Cells co-expressing hACE2-eGFP and ER-mCherry as an ER localization marker, ***a***; ssYFP-BiP and ER-mCherry-S1, b; ssYFP-BiP and ER-mCherry-S1-2, ***c***. Bar = 5 μm. (B) Lysotracker Green-stained Neuro2a cells expressing ER-mCherry as control, ER-mCherry-S1, and ER-mCherry-S1-2. Bar = 10 μm.

The concentrations of ACE2, S1, and S2 in cell lysates determined using FCS were <10 nM; thus, ACE2, S1, and S1-2 were purified and concentrated from the lysates. Western blotting using tag-specific antibodies revealed the successful recovery of intact proteins (Figure 2A). Contaminants of purified samples were confirmed by SDS-PAGE followed by silver staining (Figure 2B). However, these purified samples were used for the measurements because FCCS can specifically analyze target molecular interactions, even in the presence of non-fluorescent contaminants.

**Figure 2.**
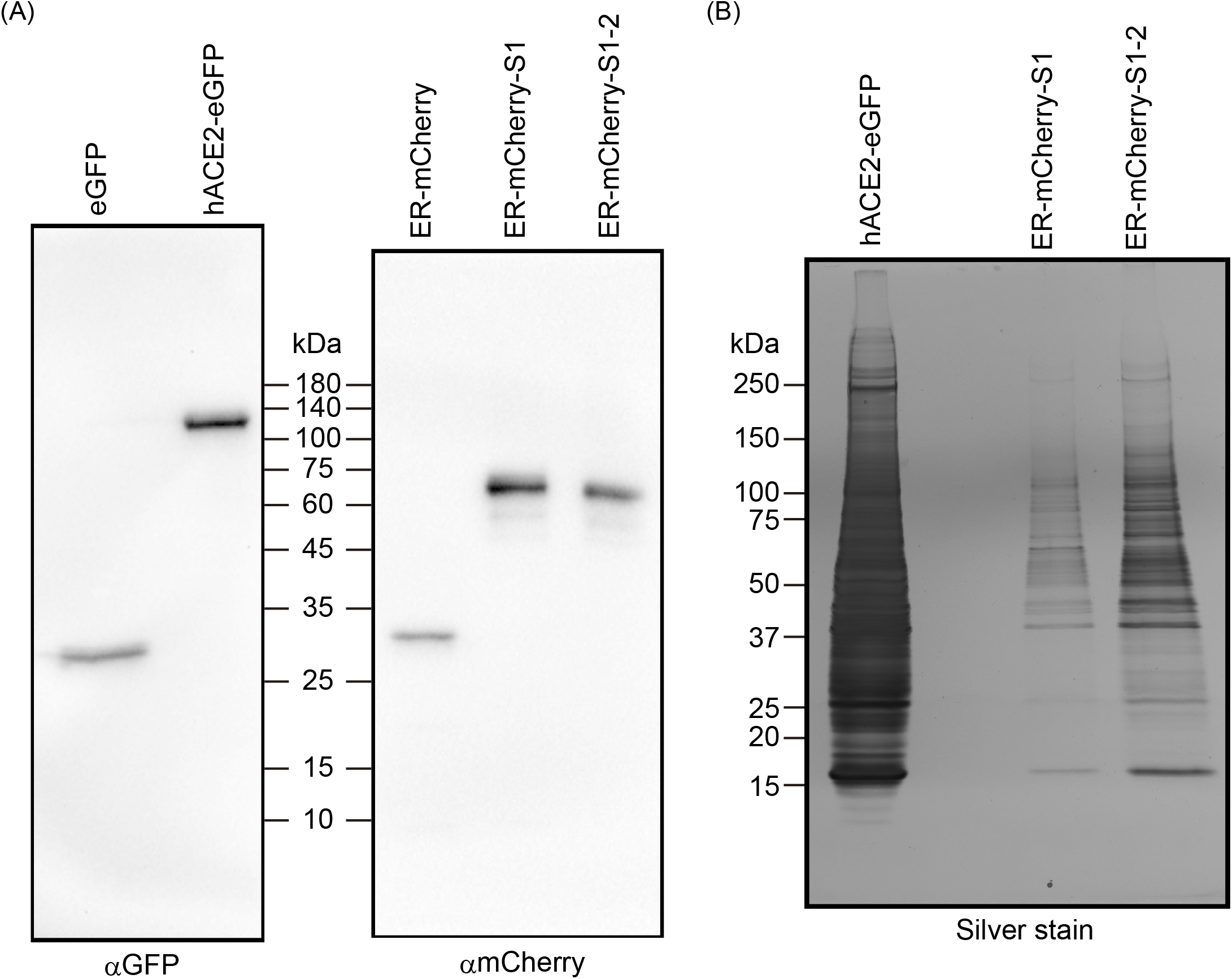
Confirmation of purified recombinant hACE2-eGFP, ER-mCherry-S1, and ER-mCherry-S1-2. (A) Western blot of purified recombinant eGFP, hACE2-eGFP, ER-mCherry, ER-mCherry-S1, and ER-mCherry-S1-2 using anti-GFP and anti-mCherry antibodies (Left and right, respectively). (B) SDS-PAGE gel followed by silver staining of purified hACE2-eGFP, ER-mCherry-S1, and ER-mCherry-S1-2.

The positive amplitude of normalized cross-correlation functions (nCCFs) indicates the interaction between two-colored fluorophores [25,29]; thus, typical nCCFs were compared when ACE2 was mixed with S1, S1-2, or mCherry monomers. The amplitude of nCCF of the mixture between ACE2 and mCherry monomers as a control was not observed (Ctrl in Figure 3A). In contrast, the amplitudes of nCCFs of the mixture between ACE2 and S1 or S1-2 were positive (green or magenta lines in Figure 3A). Furthermore, the amplitude between ACE2 and S1-2 was higher than that between ACE2 and S1 (Figure 3A). To quantitatively compare the interaction strengths among them, the relative cross-correlation amplitude (RCA), which is the ratio of the number of interacting molecules to the total number of green fluorescent molecules, was calculated. RCAs of the mixture between eGFP or mCherry monomers and ACE2, S1, or S1-2 as controls were almost zero; however, the RCAs of the mixture between ACE2 and S1 or S1-2 were significantly positive (Figure 3B). RCA in the case of S1-2 was higher than that in S1 (Figure 3B). These results suggest that both S1 and S1-2 interact with ACE2, and that S1-2 interacts more strongly than S1. Moreover, counts per molecule (CPM) values, the mean brightness of a single molecule, of eGFP- or mCherry-tagged proteins were not different from those of eGFP or mCherry monomers, suggesting that ACE2, S1, and S1-2 existed as monomers in the mixture (Figure 3C and D).

**Figure 3.**
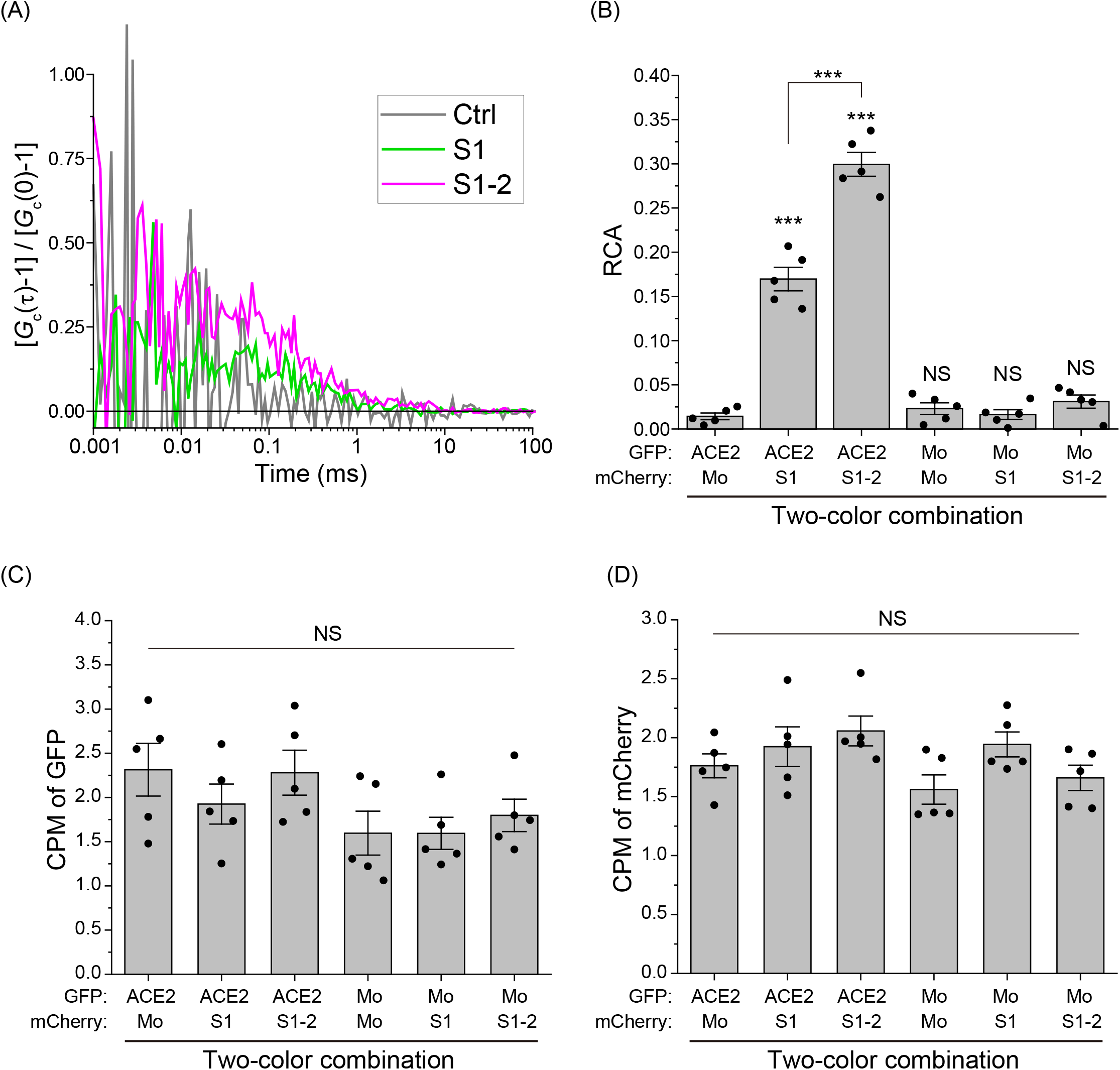
Interaction analysis between ACE2 and S1 subunit using fluorescence cross-correlation spectroscopy (FCCS). (A) Typical normalized cross-correlation functions (nCCFs: [*G*_c_(τ)-1/*G*_c_(0)-1]) of mixtures of purified mCherry monomers (Ctrl; grey), ER-mCherry-S1 (S1; green), or ER-mCherry-S1-2 (S1-2; magenta) with hACE2-eGFP. The *x*-axis shows time (τ). (B) Relative cross-correlation amplitude (RCA) of a two fluorescent color combination mixture. Mo indicates GFP or mCherry monomers. Bars indicate mean ± SE. Dots indicate independent values. Statistics: \*\*\**p* < 0.001, NS: *p* ≥ 0.05 (not significant); comparison against the mixture including hACE2-eGFP and mCherry monomers as a negative control or line-indicated paris. (C) Counts per pmolecule (CPM) of eGFP-tagged proteins during FCCS measurement. (D) Counts per pmolecule (CPM) of mCherry-tagged proteins during FCCS measurement.

## Discussion

Here, we show the interaction between ACE2 and spike protein 1 using FCCS. Our results indicate that FCCS is a powerful tool to directly and quantitatively determine the interaction between a spike protein of SARS-CoV-2 and its human receptor ACE2 in a solution that does not depend on solid-liquid phase-bases assays (e.g., pull-down assay and SPR). Solution-based measurements and a short measurement time (less than 1 min per ~10 μL sample) on FCS can be applied for high-throughput screening of inhibitory drugs and neutralization antibodies for interaction between them, likely leading to prevention of virus infection and reducing severity where infection does occur. In this report, affinity purification for a polyhistidine tag was performed, but highly purified samples could not be obtained. This is because the intracellular expression of fluorescent protein-tagged hACE2, S1, and S1-2 was low, and some proteins bound nonspecifically to the carrier for affinity purification. To obtain highly purified samples, additional affinity purification and/or other column chromatography techniques, such as gel filtration, in addition to the increased number of cells, could be used. However, highly purified samples were not required for interaction analysis using FCCS because FCCS is a fluorescence-based detection of the protein interaction with a high target specificity [22]. According to these characteristics, it could be possible to measure the interaction between purified hACE2 and S proteins in a solution mixed with a crude sample (e.g., serum addition). There are two important conditions for analyzing the interaction using FCS: (i) the specific labeling of the fluorescent tag and (ii) the expression of the full-length/intact form of the analyzed protein. We confirmed that the proteins in the samples existed in such a state using western blotting before the FCCS measurements (Figure 2A).

Spike proteins are composed of two subunits (S1 and S2). In this study, we only used the S1 subunit. CPM values obtained by FCCS analysis showed that mCherry-tagged S1 subunits exist as monomers (Figure 3C), suggesting that the S1 protein could be stable as monomers. Trimer formation of the spike protein has been reported [4,30]; however, the absence of the S2 subunit would not induce trimer formation. The monomer stability of the S1 subunit is considered advantageous for analyzing the interaction. Moreover, CPM values also showed no oligomerization of GFP-tagged and transmembrane domain-lacking hACE2 even with S1 and S1-2 (Figure 3C and D), suggesting a one-to-one interaction between ACE2 and the S1 subunit.

Regarding the subcellular localization of expressed fluorescent protein-tagged hACE2 and spike proteins, a portion of mCherry-tagged spike proteins were localized in the lysosome, although they were also observed in the ER (Figure 1). The mRFP1 and its variants can be localized in the lysosome because they are fluorescent even in an acidic environment and are resistant to proteases in the lysosome [28]. Western blotting showed no dominant fragments of mCherry-tagged proteins, indicating that lysosomal localization was not involved in the degradation of extracted proteins. Moreover, GFP-tagged hACE2 was not observed in the lysosomes because of its lower acidic stability.

Considering the current urgent medical and social requirements for the discovery of therapeutic candidates for COVID-19, including existing drugs for other diseases [31], our method for the interaction analysis between spike proteins of SARS-CoV-2 and human cell surface viral receptors using FCCS would contribute to drug screening and discovery.

## Acknowledgments

A.K. was supported by a grant from the Nakatani Foundation for novel coronavirus research, by a grant-in-aid from Hoansha Foundation, by a Japan Society for Promotion of Science (JSPS) Grant-in-Aid for Scientific Research (C) (18K06201), by a grant from the Canon Foundation, and by a grant from Hokkaido University Office for Developing Future Research Leaders (L-Station). M. K. was partially supported by a JSPS Grant-in-Aid for Scientific Research on Innovative Areas “Chemistry for Multimolecular Crowding Biosystems” (#20H04686), and a JSPS Grant-in-Aid for Scientific Research on Innovative Areas “Information physics of living matters” (#20H05522).

## Author contributions

Conceived and designed the experiments: AK. Protein purification: AF, LY, and AK. FCCS measurement: AF and AK. Microscopy: AK. Analyzed the data: AF and AK. Wrote the paper: AF, MK, and AK. All the authors agree with the publication of this paper.

## Conflicts of interest

The authors declare that they have no conflicting interests.

**Supplemental Figure Inserted cDNA sequences of hACE2-eGFP, ER-mCherry-S1 and ER-mCherry-S1-2 for protein expression.** (A) Forward sequence of hACE2-eGFP between the NheI and NotI sites. Orange: hACE2-coding region; green: eGFP-coding region; dark blue: polyhistidine tag coding region. (B and C) Forward sequences of ER-mCherry-S1 and -S1-2 between the NheI and BamHI sites. Brown: ER-sorting signal sequence-coding region; Magenta: mCherry-coding region; Cyan: S1- or S1-2-coding region; dark blue: polyhistidine tag coding region. The values on the right are the base numbers.

## Notes

### Competing Interest Statement

The authors have declared no competing interest.

