## Supplemental Figure for "Interaction between spike protein of SARS-CoV-2 and human virus receptor ACE2 using two-color fluorescence cross-correlation spectroscopy"

### Supplementary Figure

(A)

***hACE2-eGFP***

```
gctagcgttt aaacgggccc tctagaccAT GTCAAGCTCT TCCTGGCTCC TTCTCAGCCT TGTTGCTGTA ACTGCTGCTC AGTCCACCAT TGAGGAACAG 100
GCCAAGACAT TTTTGGACAA GTTTAACCAC GAAGCCGAAG ACCTGTTCTA TCAAAGTTCA CTTGCTTCTT GGAATTATAA CACCAATATT ACTGAAGAGA 200
ATGTCCAAAA CATGAATAAT GCTGGGGACA AATGGTCTGC CTTTTTAAAG GAACAGTCCA CACTTGCCCA AATGTATCCA CTACAAGAAA TTCAGAATCT 300
CACAGTCAAG CTTCAGCTGC AGGCTCTTCA GCAAAATGGG TCTTCAGTGC TCTCAGAAGA CAAGAGCAAA CGGTTGAACA CAATTCTAAA TACAATGAGC 400
ACCATCTACA GTACTGGAAA AGTTTGTAAC CCAGATAATC CACAAGAATG CTTATTACTT GAACCAGGTT TGAATGAAAT AATGGCAAAC AGTTTAGACT 500
ACAATGAGAG GCTCTGGGCT TGGGAAAGCT GGAGATCTGA GGTCGGCAAG CAGCTGAGGC CATTATATGA AGAGTATGTG GTCTTGAAAA ATGAGATGGC 600
AAGAGCAAAT CATTATGAGG ACTATGGGGA TTATTGGAGA GGAGACTATG AAGTAAATGG GGTAGATGGC TATGACTACA GCCGCGGCCA GTTGATTGAA 700
GATGTGGAAC ATACCTTTGA AGAGATTAAA CCATTATATG AACATCTTCA TGCCTATGTG AGGGCAAAGT TGATGAATGC CTATCCTTCC TATATCAGTC 800
CAATTGGATG CCTCCCTGCT CATTTGCTTG GTGATATGTG GGGTAGATTT TGGACAAATC TGTACTCTTT GACAGTTCCC TTTGGACAGA AACCAAACAT 900
AGATGTTACT GATGCAATGG TGGACCAGGC CTGGGATGCA CAGAGAATAT TCAAGGAGGC CGAGAAGTTC TTTGTATCTG TTGGTCTTCC TAATATGACT 1000
CAAGGATTCT GGGAAAATTC CATGCTAACG GACCCAGGAA ATGTTCAGAA AGCAGTCTGC CATCCCACAG CTTGGGACCT GGGGAAGGGC GACTTCAGGA 1100
TCCTTATGTG CACAAAGGTG ACAATGGACG ACTTCCTGAC AGCTCATCAT GAGATGGGGC ATATCCAGTA TGATATGGCA TATGCTGCAC AACCTTTTCT 1200
GCTAAGAAAT GGAGCTAATG AAGGATTCCA TGAAGCTGTT GGGGAAATCA TGTCACTTTC TGCAGCCACA CCTAAGCATT TAAAATCCAT TGGTCTTCTG 1300
TCACCCGATT TTCAAGAAGA CAATGAAACA GAAATAAACT TCCTGCTCAA ACAAGCACTC ACGATTGTTG GGACTCTGCC ATTTACTTAC ATGTTAGAGA 1400
AGTGGAGGTG GATGGTCTTT AAAGGGGAAA TTCCCAAAGA CCAGTGGATG AAAAAGTGGT GGGAGATGAA GCGAGAGATA GTTGGGGTGG TGGAACCTGT 1500
GCCCCATGAT GAAACATACT GTGACCCCGC ATCTCTGTTC CATGTTTCTA ATGATTACTC ATTCATTCGA TATTACACAA GGACCCTTTA CCAATTCCAG 1600
TTTCAAGAAG CACTTTGTCA AGCAGCTAAA CATGAAGGCC CTCTGCACAA ATGTGACATC TCAAACTCTA CAGAAGCTGG ACAGAAACTG TTCAATATGC 1700
TGAGGCTTGG AAAATCAGAA CCCTGGACCC TAGCATTGGA AAATGTTGTA GGAGCAAAGA ACATGAATGT AAGGCCACTG CTCAACTACT TTGAGCCCTT 1800
ATTTACCTGG CTGAAAGACC AGAACAAGAA TTCTTTTGTG GGATGGAGTA CCGACTGGAG TCCATATGCA GACGGACCGG TCGCCACCAT GGTGAGCAAG 1900
GGCGAGGAGC TGTTCACCGG GGTGGTGCCC ATCCTGGTCG AGCTGGACGG CGACGTAAAC GGCCACAAGT TCAGCGTGTC CGGCGAGGGC GAGGGCGATG 2000
CCACCTACGG CAAGCTGACC CTGAAGTTCA TCTGCACCAC CGGCAAGCTG CCCGTGCCCT GGCCCACCCT CGTGACCACC CTGACCTACG GCGTGCAGTG 2100
CTTCAGCCGC TACCCCGACC ACATGAAGCA GCACGACTTC TTCAAGTCCG CCATGCCCGA AGGCTACGTC CAGGAGCGCA CCATCTTCTT CAAGGACGAC 2200
GGCAACTACA AGACCCGCGC CGAGGTGAAG TTCGAGGGCG ACACCCTGGT GAACCGCATC GAGCTGAAGG GCATCGACTT CAAGGAGGAC GGCAACATCC 2300
TGGGGCACAA GCTGGAGTAC AACTACAACA GCCACAACGT CTATATCATG GCCGACAAGC AGAAGAACGG CATCAAGGTG AACTTCAAGA TCCGCCACAA 2400
CATCGAGGAC GGCAGCGTGC AGCTCGCCGA CCACTACCAG CAGAACACCC CCATCGGCGA CGGCCCCGTG CTGCTGCCCG ACAACCACTA CCTGAGCACC 2500
CAGTCCAAGC TGAGCAAAGA CCCCAACGAG AAGCGCGATC ACATGGTCCT GCTGGAGTTC GTGACCGCCG CCGGGATCAC TCTCGGCATG GACGAGCTGT 2600
ACAGTCACCA TCATCACCAT CACTAAagcg gccgc
```

(B)

***ER-mCherry-S1***

```
gctagcATGC TGCTATCCGT GCCGTTGCTG CTCGGCCTCC TCGGCCTGGC CGTCGCCGGA CCGGTCGCCA CCATGGTGAG CAAGGGCGAG GAGGACCAGA 100
TGGCCATCAT CAAGGAGTTC ATGCGCTTCA AGGTGCACAT GGAGGGCTCC GTGAACGGCC ACGAGTTCGA GATCGAGGGC GAGGGCGAGG GCCGCCCCTA 200
CGAGGGCACC CAGACCGCCA AGCTGAAGGT GACCAAGGGT GGCCCCCTGC CCTTCGCCTG GGACATCCTG TCCCCTCAGT TCATGTACGG CTCCAAGGCC 300
TACGTGAAGC ACCCCGCCGA CATCCCCGAC TACTTGAAGC TGTCCTTCCC CGAGGGCTTC AAGTGGGAGC GCGTGATGAA CTTCGAGGAC GGCGGCGTGG 400
TGACCGTGAC CCAGGACTCC TCCCTGCAGG ACGGCGAGTT CATCTACAAG GTGAAGCTGC GCGGCACCAA CTTCCCCTCC GACGGCCCCG TAATGCAGAA 500
GAAGACCATG GGCTGGGAGG CCTCCTCCGA GCGGATGTAC CCCGAGGACG GCGCCCTGAA GGGCGAGATC AAGCAGAGGC TGAAGCTGAA GGACGGCGGC 600
CACTACGACG CTGAGGTCAA GACCACCTAC AAGGCCAAGA AGCCCGTGCA GCTGCCCGGC GCCTACAACG TCAACATCAA GTTGGACATC ACCTCCCACA 700
ACGAGGACTA CACCATCGTG GAACAGTACG AACGCGCCGA GGGCCGCCAC TCCACCGGCG GCATGGACGA GCTGTACAAG TCCGGACTCA GATCTCGAGC 800
TAGCTTTGAG ATTGACAAAG GAATTTACCA GACCTCTAAT TTCAGGGTTG TTCCCTCAGG AGATGTTGTG AGATTCCCTA ATATTACAAA CTTGTGTCCT 900
TTTGGAGAGG TTTTTAATGC TACTAAATTC CCTTCTGTCT ATGCATGGGA GAGAAAAAAA ATTTCTAATT GTGTTGCTGA TTACTCTGTG CTCTACAACT 1000
CAACATTTTT TTCAACCTTT AAGTGCTATG GCGTTTCTGC CACTAAGTTG AATGATCTTT GCTTCTCCAA TGTCTATGCA GATTCTTTTG TAGTCAAGGG 1100
AGATGATGTA AGACAAATAG CGCCAGGACA AACTGGTGTT ATTGCTGATT ATAATTATAA ATTGCCAGAT GATTTCATGG GTTGTGTCCT TGCTTGGAAT 1200
ACTAGGAACA TTGATGCTAC TTCAACTGGT AATTATAATT ATAAATATAG GTATCTTAGA CATGGCAAGC TTAGGCCCTT TGAGAGAGAC ATATCTAATG 1300
TGCCTTTCTC CCCTGATGGC AAACCTTGCA CCCCACCTGC TCTTAATTGT TATTGGCCAT TAAATGATTA TGGTTTTTAC ACCACTACTG GCATTGGCTA 1400
CCAACCTTAC AGAGTTGTAG TACTTTCTTT TGAACTTTTA AATGCACCGG CCACGGTTTG TGGACCAAAA TTATCCACTG ACCTTATTAA GAACCAGTGT 1500
GTCAATTTTA ATTTTAATGG ACTCACTGGT ACTGGTGTGT TAACTCCTTC TTCAAAGAGA TTTCAACCAT TTCAACAATT TGGCCGTGAT GTTTCTGATT 1600
TCACTGATTC CCACCATCAT CACCATCACT AAggatcc
```

(C)

***ER-mCherry-S1-2***

```
gctagcATGC TGCTATCCGT GCCGTTGCTG CTCGGCCTCC TCGGCCTGGC CGTCGCCGGA CCGGTCGCCA CCATGGTGAG CAAGGGCGAG GAGGACCAGA 100
TGGCCATCAT CAAGGAGTTC ATGCGCTTCA AGGTGCACAT GGAGGGCTCC GTGAACGGCC ACGAGTTCGA GATCGAGGGC GAGGGCGAGG GCCGCCCCTA 200
CGAGGGCACC CAGACCGCCA AGCTGAAGGT GACCAAGGGT GGCCCCCTGC CCTTCGCCTG GGACATCCTG TCCCCTCAGT TCATGTACGG CTCCAAGGCC 300
TACGTGAAGC ACCCCGCCGA CATCCCCGAC TACTTGAAGC TGTCCTTCCC CGAGGGCTTC AAGTGGGAGC GCGTGATGAA CTTCGAGGAC GGCGGCGTGG 400
TGACCGTGAC CCAGGACTCC TCCCTGCAGG ACGGCGAGTT CATCTACAAG GTGAAGCTGC GCGGCACCAA CTTCCCCTCC GACGGCCCCG TAATGCAGAA 500
GAAGACCATG GGCTGGGAGG CCTCCTCCGA GCGGATGTAC CCCGAGGACG GCGCCCTGAA GGGCGAGATC AAGCAGAGGC TGAAGCTGAA GGACGGCGGC 600
CACTACGACG CTGAGGTCAA GACCACCTAC AAGGCCAAGA AGCCCGTGCA GCTGCCCGGC GCCTACAACG TCAACATCAA GTTGGACATC ACCTCCCACA 700
ACGAGGACTA CACCATCGTG GAACAGTACG AACGCGCCGA GGGCCGCCAC TCCACCGGCG GCATGGACGA GCTGTACAAG TCCGGACTCA GATCTCGAGC 800
TTCCTTCACT GTAGAAAAAG GAATCTATCA AACTTCTAAC TTTAGAGTCC AACCAACAGA ATCTATTGTT AGATTTCCTA ATATTACAAA CTTGTGCCCT 900
TTTGGTGAAG TTTTTAACGC CACCAGATTT GCATCTGTTT ATGCTTGGAA CAGGAAGAGA ATCAGCAACT GTGTTGCTGA TTATTCTGTC CTATATAATT 1000
CCGCATCATT TTCCACTTTT AAGTGTTATG GAGTGTCTCC TACTAAATTA AATGATCTCT GCTTTACTAA TGTCTATGCA GATTCATTTG TAATTAGAGG 1100
TGATGAAGTC AGACAAATCG CTCCAGGGCA AACTGGAAAG ATTGCTGATT ATAATTATAA ATTACCAGAT GATTTTACAG GCTGCGTTAT AGCTTGGAAT 1200
TCTAACAATC TTGATTCTAA GGTTGGTGGT AATTATAATT ACCTGTATAG ATTGTTTAGG AAGTCTAATC TCAAACCTTT TGAGAGAGAT ATTTCAACTG 1300
AAATCTATCA GGCCGGTAGC ACACCTTGTA ATGGTGTTGA AGGTTTTAAT TGTTACTTTC CTTTACAATC ATATGGTTTC CAACCCACTA ATGGTGTTGG 1400
TTACCAACCA TACAGAGTAG TAGTACTTTC TTTTGAACTT CTACATGCAC CAGCAACTGT TTGTGGACCT AAAAAGTCTA CTAATTTGGT TAAAAACAAA 1500
TGTGTCAATT TCAACTTCAA TGGTTTAACA GGCACAGGTG TTCTTACTGA GTCTAACAAA AAGTTTCTGC CTTTCCAACA ATTTGGCAGA GACATTGCTG 1600
ACACTACTGA TGCTCACCAT CATCACCATC ACTAAggatc c
```
